# Effects of simulated acid rain on soil respiration rate and soil bacterial diversity in a *Phyllostachyspubescens* forest in subtropical China

**DOI:** 10.1101/688838

**Authors:** Nan Wang, Xiaocheng Pan

## Abstract

Acid rain has been regarded as a global environmental concern due to its negative effects on global ecosystems. In this study, we investigated the effects of simulated acid rain (SAR) on soil respiration rate and soil bacterial diversity in a Moso bamboo (*phyllostachyspubescens*) forest in subtropical China. Experimental results showed a similar seasonal pattern of soil respiration rates underdifferent SAR treatments. Seasonal mean soil respiration rates for CK (control, deionized water, pH 6.7), T1 (pH 5.6), T2 (pH 4.0) and T3 (pH 2.5) treatments were 3.44, 4.80, 4.35 and 4.51 μ mol m^−2^ s^−1^, respectively. One-way analysis of variance indicated that the SAR exposure had no significant effect on soil respiration (p>0.1) and soil microbial biomass (p>0.1). Soil bacterial community diversity was calculated as the Shannon-Wiener diversity index and the results showed that only T3 treatment had significant effects on soil bacterial diversity. The DGGE analysis results revealed that T1 and CK soils had closer association and were related to the T2 soil, while T3 soil was distinctly different from the other treatments. This work highlights that the effects of SAR are important to consider in assessing the soil respiration rate, particularly under the scenario of increasing acid rain pollution.

## 1. Introduction

Recently, with the rapid growth of worldwide economy and urban population, more and more sulfur dioxide (SO_2_) and nitrogen oxides (NO_x_) gases are produced during the combustion of fossil fuels within thermal power plants and automobiles. The emission of these gases into atmosphere could mix with water vapor in the air to form sulfuric and nitric acid, which later falls as acid rain [1, 2]. Acid rain has become a global environmental problem and received worldwide attention due to its environmental damage, including the acidification of soil [3], decrease of microbial community function and enzyme activities [4–6], negative effects on vegetation, in particular forests [7], and the changes of soil species composition [8].

In China, particularly in the southern part, acid rain is also a serious environmental hazard as most of the soil there is acidic. South China, Europe and North America have become the three most severely affected regions by acid rain in the world [9, 10]. As one of the largest CO_2_ fluxes in the global carbon (C) cycle, soil respiration contributes to 68-98 Pg-C to the atmosphere annually [11, 12]. Moreover, belowground Caccounts for more than two-thirds of the terrestrial C stock, and roots and microorganisms in the soils play an important role in atmospheric CO_2_ respiration. Thus, deep understanding of soil respiration process will help us expand the knowledge of the terrestrial C cycle [13].

During the past decades, researchers mainly focused on how acid rain impacted the soil respiration in forest ecosystems. Acid deposition might directly affect soil respiration by changing microbial activity, enzyme activity, and the composition of the microbial population, as stated above. However, the results obtained in previous study were inconsistent. Chen et al. [9] found little impact on the soil respiration by simulated acid rain (SAR), while Blagodaskaya and Anderson [14] revealed that soil respiration corresponded with SAR loads, which might be associated with the adaptability of bacterial community [15]. Likewise, acid deposition might indirectly influence soil respiration due to the decrease of pH. Spain [16] revealed that organic C content in soil increased when pH in soil decreased, while Baath and Anderson [17] observed an increase of soil respiration rates along with the decrease of pH in soil.

*Phyllostachyspubescens*, a typical and economic bamboo species, is widely grown in South China, due to its rapid growth rate and forest formation, high revenue, widespread use, and high regeneration capacity. In China, the bamboo-growing area is increasing year by year and *phyllostachyspubescens* (referred as bamboo forest afterwards) forest has become an important ecosystem type. Unfortunately, few investigations have focused on the impacts of acid rain on soil respiration in bamboo forest, and to our knowledge, the long-term *in situ* experiment was missing. Also, the information about the effect of acid rain on bacteria microbial community in bamboo forest is rare. Thus, studying soil respiration and microbial community in bamboo forest (subtropical forest) subject to SAR treatments is important for understanding their functions in C cycling/ecosystem C flux.

In this study, we measured the soil respiration, soil microbial biomass (characterized as soil microbial biomass C (C_mic_) and soil microbial biomass nitrogen (N_mic_)) and microbial community change in a bamboo forest, a subtropical soil environment. The selected forest was subject to 10 months of artificial acid rain to evaluate if soil respiration and the microbial community is altered by different SAR levels.

## 2. Materials and Methods

### 2.1 Site description

In 2015, experiments were conducted at Tianmu Mountain (30.30°N, 119.45°E) near Hangzhou city, in Zhejiang province, China. The detailed information about Tianmu Mountain could be seen in Wang et al. [18]. The bamboo forest is at 500 m elevation and adjoins evergreen broad-leaf forest and commonly mixed grow with *Castanopsissclerophylla*, *Castanopsismysinaefolia*, *Zelkovaschneideriana*, and *Liquidambar formosana*. The area of bamboo forest occupies 875,000m^2^, and understory species are rare, including *Camellia sinensis*, *Euryahebeclados*, *Cyclobalanopsisgracilis*, *Cyclobalanopsismyrsinifolia*, *Rhododendron ovatum* and *Lithocarpusbrevicaudatus*. The properties of soil at the end of the experiment are shown in Table 1.

### 2.2 SAR treatment

The experiment was conducted since September, 2016. Twelve sample plots (10 m×10 m) were selected and divided into four groups, and the twelve sample plots were almost at the same elevation to avoid the effect of mountain slope gradient on soil respiration as much as possible. According to the acid rain characteristic and acid deposition levels in Lin’ An, Jiangsu province, four groups of experiments were designed as follows: control experiment (termed as CK), only deionized water was applied to the experimental sites and the pH was approximately 6.7; T1-T3 experiment, prepared acid rain was applied to the corresponding sites and the pH was 5.6, 4.0 and 2.5 respectively. The simulated acid rain was prepared from deionized water and contained H_2_SO_4_ and HNO_3_ with a mole ratio of 4.5:1 [19]. Throughout the duration of the experiment, 10 L of the simulated acid rain were applied to each site twice a week (or postponed in case of rain or high soil humidity). In order to ensure the acid rain permeated into the soil evenly, we used a simulation apparatus capable of delivering droplet sizes in the range of 1.0 to 1.2 mm diameter. The experimental sites were subjected to simulated acid rain treatments for 10 months.

### 2.3 Soil respiration rate measurement

Soil respiration rate was measured according to Chen et al. [1]. The PVC soil collar (20 cm in diameter) was permanently installed (5 cm) into the soil in each site and soil packing by PVC collar was minimized. In order to minimize the impact of aboveground respiration by living plants during soil respiration rate measurement, we removed the living plants within the soil collar completely prior to measurements. Measurements were generally implemented once a month from September 2016 to July 2017. The detailed information about soil temperature and moisture measurement can be found in Chen et al. [1]. Each measurement started at 09:00 am and the whole process lasted for about 2-3 hour, including 30 min for preheat, and transport [20]. In order to guarantee the accuracy, a preheat measurement was performed to exclude the residual gas inside the instrument prior to formal measurement. Furthermore, during each measurement, the distance between surveyors and gas analyzer should be more than 2 m to avoid the disturbance. The moist soil samples at each site were collected and pretreated for C_mic_ and N_mic_ analysis [21]. In brief, The C_mic_ and N_mic_ values were determined on the <2-mm mesh field-moist samples. Soil C_mic_ was estimated on a 7.3-g oven-dry equivalent of field-moist soil sample by the chloroform-fumigation-extraction method and soil N_mic_ was determined by chloroform-fumigation-incubation method using a 7.3-g oven-dry equivalent of field-moist soil sample, after adjusting the moisture content to 55%. The correction factors applied to C_mic_ and N_mic_ calculation were 0.45 and 0.57. Soil temperature (°C) and moisture (g water/kg soil) at the depth of 5 cm were monitored adjacent to each PVC collar using a probe connected to the Li-8100 during the soil respiration rate measurements.

### 2.4 Statistical analysis

Soil respiration rate in each treatment was calculated as the mean of the measurements from 3 collars. The significant level of the soil respiration rate among the SAR treatments (including CK) was tested using a *t*-test proposed by Luo et al. [42]. One-way ANOVA was used to test the SAR effects on bacterial diversity and microbial biomass. All the statistical analyses were performed using Excel 2007 (Microsoft Inc. Seattle, WA, USA) and the SPSS software version 11.0.

### 2.5 Bacterial community

At the end of the experiments, soils in twelve sample plots were collected to analyze bacterial microbial community and ten cores (5 cm diameter × 20 cm length) were taken from each sampling plot and mixed. The twelve samples were shipped to lab as soon as possible and subsequently sieved through a 2 mm mesh to avoid the interference of plant debris and soil fauna. Total DNA were firstly extracted according to Zhou et al. [22] and then purified as per Cahyani et al. [23]. The DNA yield, quality and purity were assessed as described by Chang et al. [24]. The universal bacterial primers, PRBA338f and PRUN518r, located at the V3 region of the 16S rRNA genes of bacterioplankton, were used to amplify the variable V3 region of 16S rDNA. The detailed procedure for PCR amplication and DGGE analysis was implemented as described previously by Chang et al. [24]. Bacterial community diversity was calculated as the Shannon-Wiener diversity index [22].

## 3. Results and Discussion

### 3.1 Effect of SAR on soil respiration rates

Soil temperature was measured periodically during the whole experiment and the measured results showed that the temperature in soil seasonally changed accompanying with the change of air temperature in experimental site (**Table 2**). Soil temperature and moisture are generally considered two basic factors in controlling soil respiration process. In our study, when the entire experimental period was considered, seasonal variability of soil respiration rate was mainly controlled by soil temperature, as the absolute value of difference in moisture content among all the sampled soil was small. The moisture content probably mediated the responses of soil respiration rate to temperature [43].

**Figure 1** showed the seasonal variation of soil respiration rates subject to SAR exposure. For each group experiment, the soil respiration rates varied seasonally following a similar trend with soil temperature: from September 2016 to January 2017, soil respiration rate decreased during winter, followed by a sharp increase during spring and summer from March 2017 to July 2017. This observation was in accordance with that reported by Chen et al. [1]. Seasonal mean soil respiration rate for the CK, T1, T2 and T3 treatments were 3.44, 4.80, 4.35 and 4.51 μmol m^−2^ s^−1^, respectively (**Figure 2**). However, the soil respiration rates among the four group experiments were not regular. For example, the soil respiration rate of T2 was lowest at the beginning of the experiment while it reached to highest value at the end of the experiment. At each sampling point, no typical trend with the strength of SAR was observed, which suggested that it’ s not easy to clearly state how the different SAR treatments impacted on the soil respiration rates herein.

**Figure 1.**
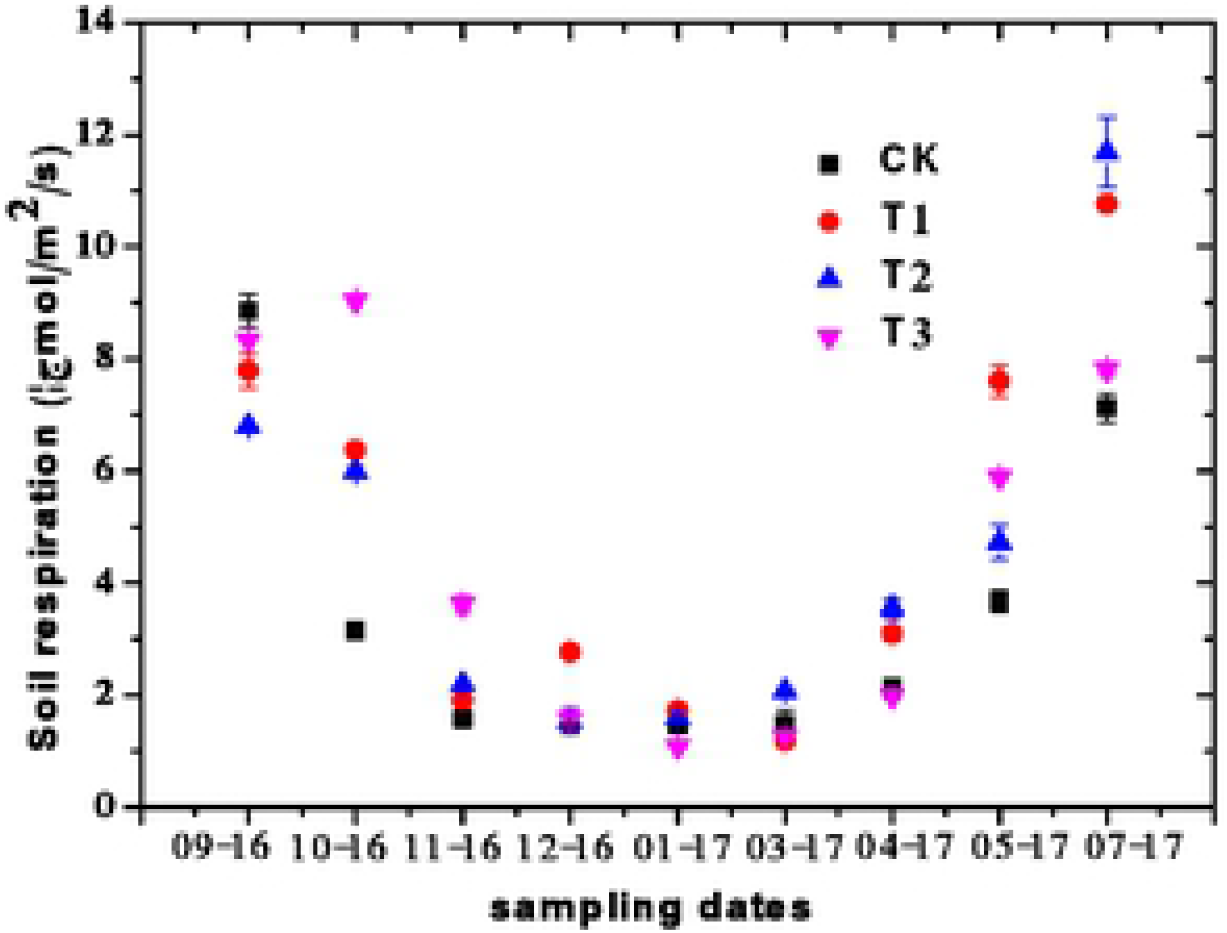
Seasonal variations of soil respiration under different SAR treatments (sampling date: month year)

**Figure 2.**
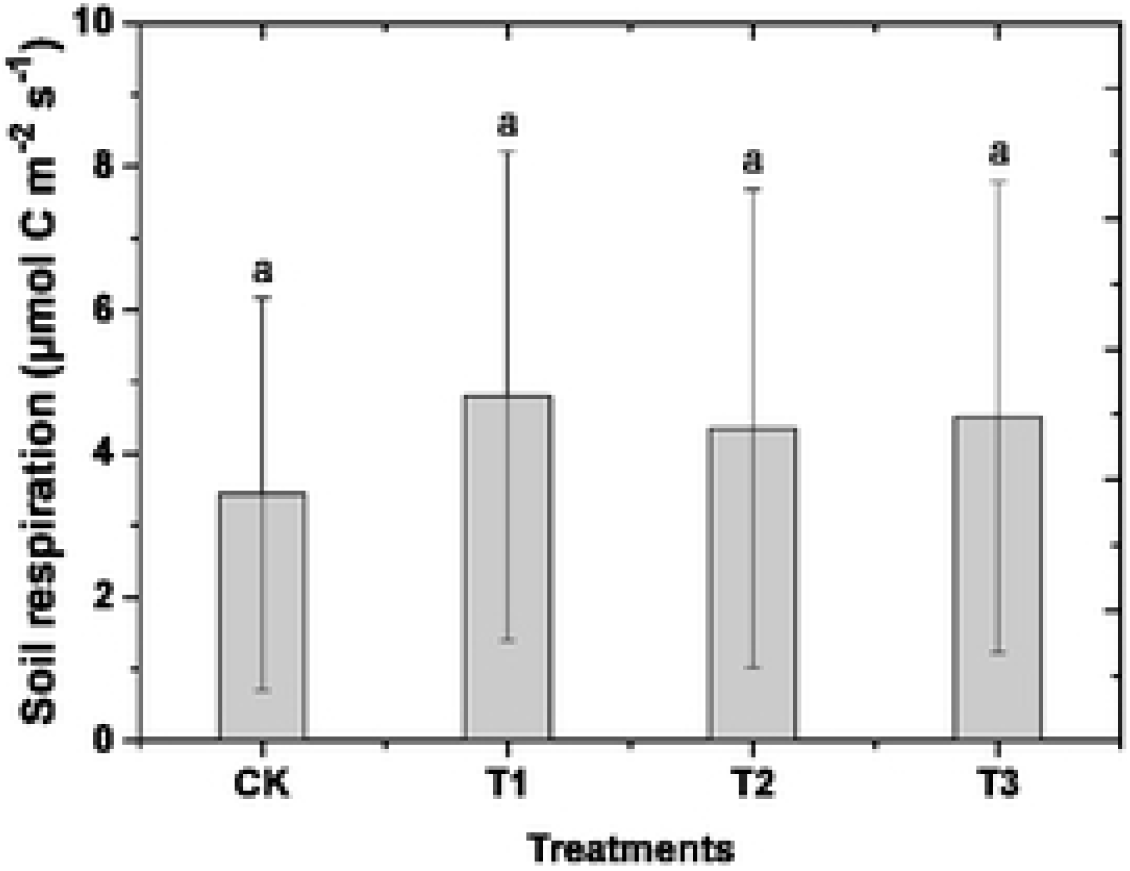
Mean soil respiration rates for different SAR treatments (Error bars are standard error of the mean; the same letter above the column are not significantly different between different treatments)

Compared with CK (control, deionized water, pH 6.7), all the SAR treatments in this study (including T1 (pH 5.6), T2 (pH 4.0), T3 (pH 2.5)) induced a positive effect on the soil respiration rate. Soil respiration rates under T1, T2 and T3 treatments were enhanced by 39.5%, 26.5% and 31.1%, respectively, relative to that of CK treatment. However, ANOVA resultsindicated that different SAR exposure of T1, T2 and T3 had no significant effects on soil respiration (p>0.1), although SAR are generally considered to have inhibition effect on soil respiration rate [40]. Although effect of SAR treatments on soil respiration has been widely studied, results from these studies were inconsistent. For example, Will et al. [25] revealed that acid rain impacted little on the soil CO_2_ flux while Zelles et al. [26] observed a decreased CO_2_ emission when an artificial acidic soil subject to SAR exposure and this negative effects might be contributed to the inhibited activity of soil microbes by low pH in the soil. What’s more, an enhanced soil CO_2_ flux was also reported when a low concentration of simulated acid rain was applied to the soil [27], which was in consistent with our results in this study. Organisms in soil may mediate themselves to the changing acid environment and meanwhile soil could buff the SAR effects on soil to a certain degree. The enhanced effect may result from nutrients, such as nitrate nitrogen (NO_3_^−^-N), that are used to acidify the simulated acid rain [41]. The inconsistent results with regard to the effect of SAR on soil respiration could be justified using several lines of reasoning, including the length of experimental period [1], and the applied simulated acid rain treatment[28]. The soil respiration rate almost maintained constant at pH 4 and pH 6, while decreased 20% at pH 3. Also, the effect of SAR on soil respiration rate was reported to be relied on geographic locations [28]. Compared with microbes in temperate soil, SAR treatment impacted the microbes in subarctic soil more greatly. And soil respiration rates are more easily to be affected by SAR in dry nutrient-poor forests than in medium and mesic forests.

Soil enzyme activity is the direct expression of the soil microbial community to the metabolic requirements and available [9]. The soil enzymatic activities, including phosphatase, urease and sucrasewere shown in **Table 1**. SAR treatments (T1 and T2) resulted in a decrease in urease and sucrase activity, although the differences between CK and SAR treatments were not significant. This finding might be attributed to a potential trade-off between the organisms showing positive responses to SAR and others showing negative responses.

### 3.2 Effect of SAR on soil microbial biomass

**Figure 3** showed the variations of soil microbial biomass subject to SAR exposure and the average values were plotted in the figure. The soil microbial biomass of CK treatment in September, 2014 was not included in the figure as the soil samples were lost during transportation.

**Figure 3.**
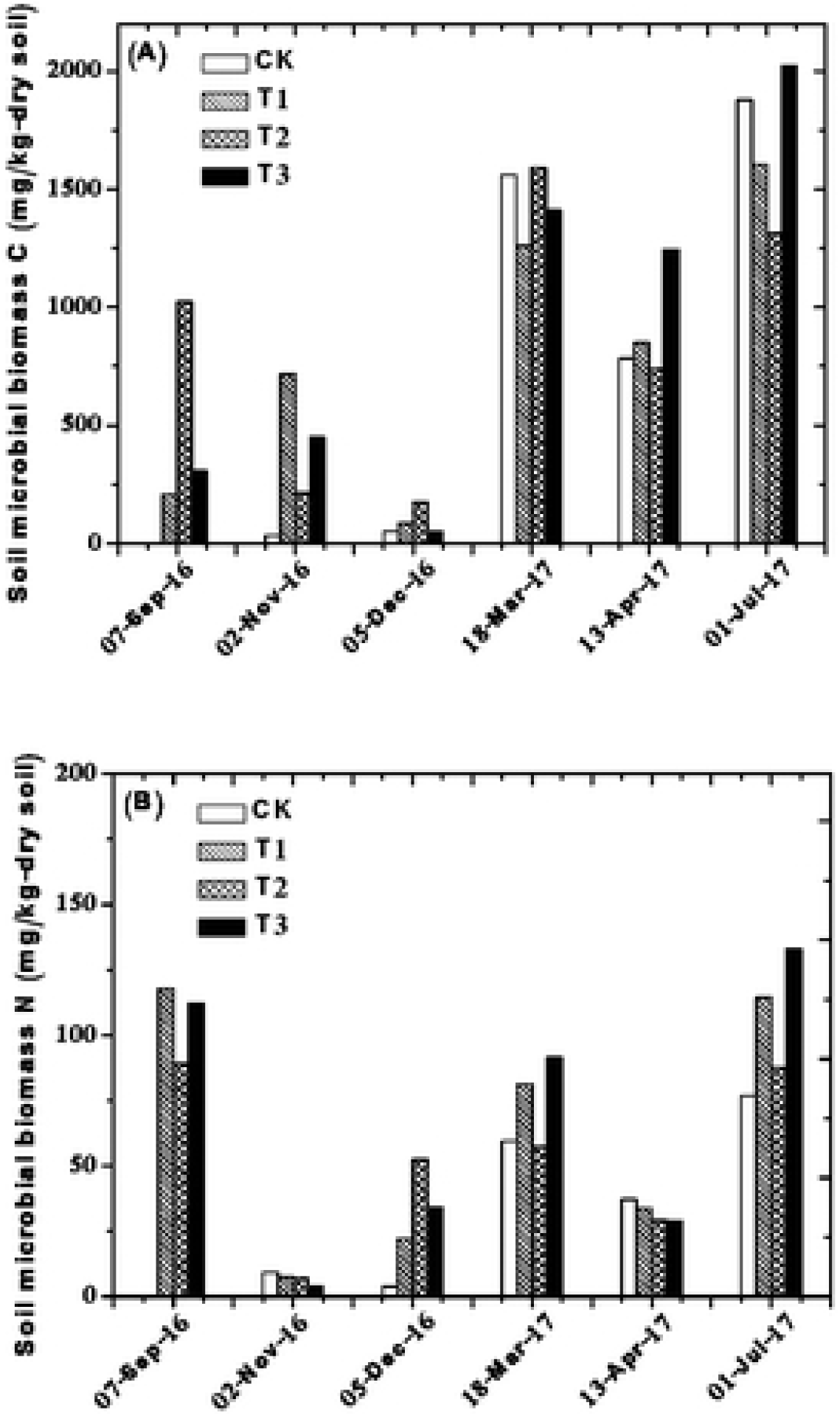
Seasonal variations of soil microbial biomass C (A) and N (B) under different SAR treatments

In winter (November/December, 2016), the concentrations of Cmic and Nmic were generally lower than that in the other sample dates. Possible explanation for the observation would be the lower microbial activity of the soil in the winter [46]. It was observed a more pronounced temporal fluctuation of Nmic values compared with those of Cmic, in agreement with those reported by Moore et al. [21]. Although C and N are the main compositions of microorganisms, their concentrations in microbes vary greatly, especially the N content, and notably depend on the growth stage of microbes. The soil pH held a clearly effect on the microbial biomass and low soil pH would lead to a low microbial biomass content [29]. However, ANOVA analysis results indicated that the difference in soil microbial biomass subject to SAR exposure was not significant (p>0.1) in this study. Furthermore, the concentrations of Cmic and Nmic at all the SAR treatments were comparable or even stimulated compared with CK treatment, which was in consistent with the trend of soil respiration rate. The microbial community might have adapted to the simulated acid rain conditions during the long-term experiment. Neither Cmic nor Nmicwas significantly correlated with soil pH. Possible explanations for this observation were that the pH of soils used in this study was acidic, and a small change of pH caused by acid rain had a minimum effect on the soil microbial biomass. Carter and Rennie[30] revealed that higher pH would enhance microbial biomass content and microbial biomass content would conversely get lost with a lower pH applied. Besides the soil pH, the microbial biomass content would also be affected by other environmental conditions. Wolters [31] found that the decreased soil pH did not impact microbial biomass content until the soil pH was lower than 2 or 3. Besides, the type of acid rain (sulfuric acid or nitric acid) would influence the microbial biomass content. Nitric acid has proven to be capable of positively and negatively impacting microbial biomass, and the variable effects mainly depend on the soil’s N threshold value. For example, Bewley and Stotzky[32] found an inhibitory effect of nitric acid on microbial biomass and activity; while Killham et al. [33] reported that nitric acid showed a stimulated effect on microbial biomass and activity exposure to nitric acid.

### 3.3 Effect of SAR on soil bacterial community diversity

The bacterial community diversity was presented as Shannon index. As shown in **Table 3**, only soil bacterial diversity of T3 was significantly different from that of CK based on the obtained value of LSD0.05. Cluster analysis showed that T1 and CK soils had closer association and were related to the T2 soil, while T3 soil was distinctly different from the other treatments (**Figure 4**). The result clearly demonstrated that DGGE profiles revealed marked differences in the response of soil bacterial communities under different SAR treatments.

**Figure 4.**
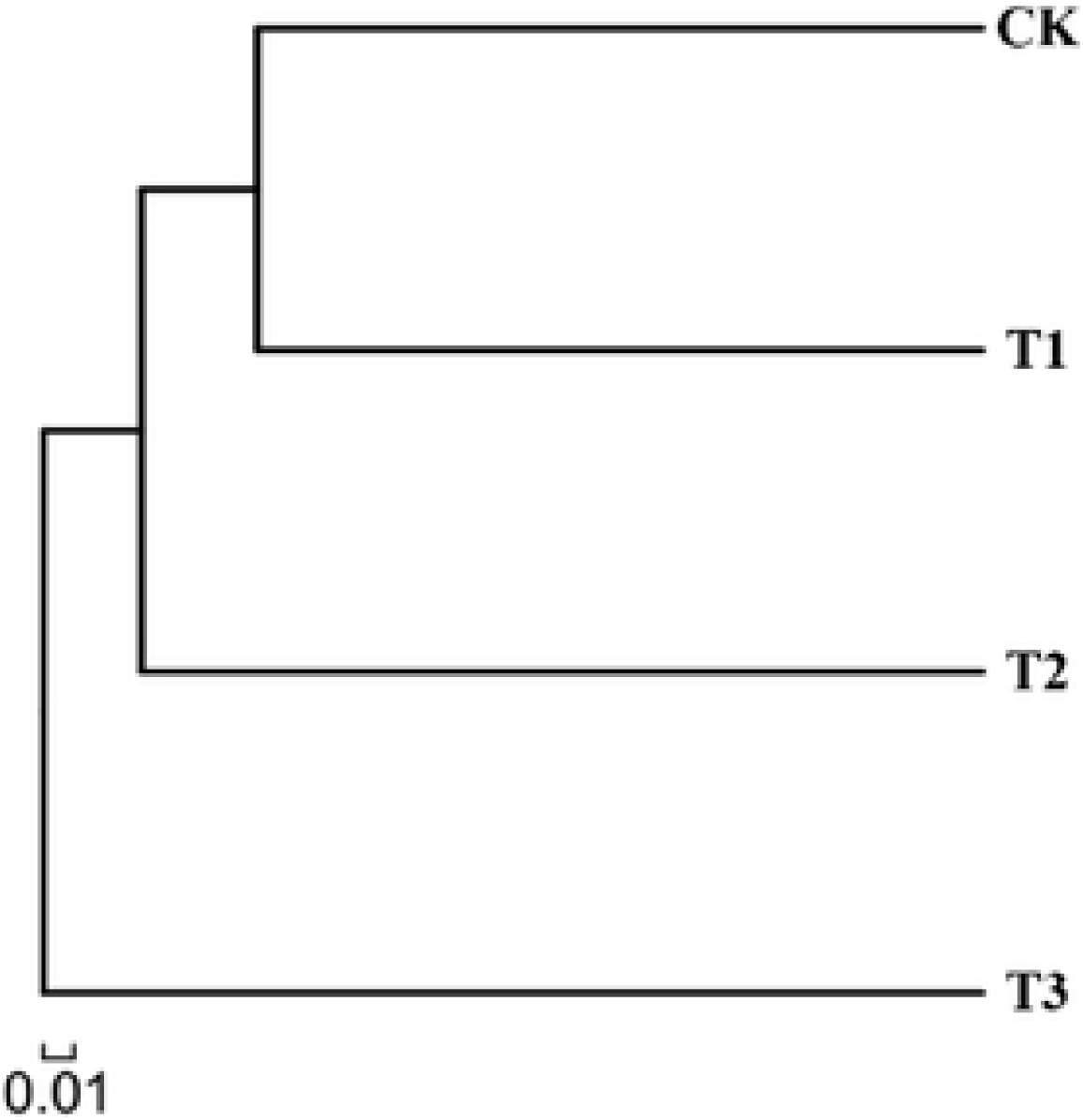
Cluster analysis of the DGGE profiles of 16S rDNA amplified from the tested soils

Soil bacterial microbial diversity may be influenced by simulated acid rainexposure [44, 45]. However, the obtained findings with regard to the effect of SAR on soil bacterial microbial diversity seemed controversial. For example, Pennanen et al. [34] reported that the amount of bacterial measured decreased with increasing pH and bacteria were more affected than fungi by the acidification. Anderson and Domsch[35] also found that the total microbial biomass was more sensitive to acidic pH than to a neutral pH. Besides, McColl and Firestone [36] and Stemmer et al. [37] observed that acid rain hardly impacted bacterial community in soil. In contrast, Pennanen et al. [38] showed that the bacteria in soil microbial community would increase when humus pH was decreased and the bacteria in soil could adapt to the new acidic environment step by step. Similarly, Wang et al. [39] found a stimulated effect on soil microbial diversity after being subjected to simulated acid rain (pH 4.5/5.5). In our study, a high acid not only stimulated soil respiration (compared with CK experiment) but also increased soil bacterial community diversity. Up to now there was no consensus in this argument and many factors could contribute to this contradiction, like the treatment period, species, and the soil sampling procedure used, which deserves further research in future. In present study, bacterial community was analyzed by DGGE method with universal bacterial primers, which could reflect the genetic diversity of a microbial community and has the advantages of being reliable, reproducible, rapid, and allows screening of multiple samples. However, it also has several limits, such as effect of variable DNA extraction efficiency on DGGE profiles, similarities in the mobility characteristics of the polyacrylamide gel of DNA fragments with different sequences, and dependence of DGGE profiles on soil type being tested and choice of primers [24]. Therefore, different approaches such as PLFA analysis, Biolog and molecular technology should be used to better investigate the effect of acid rain on soil microbial communities.

## 4. Conclusions

It can be concluded that the different SAR treatments showed similar seasonal pattern of soil respiration rate in *phyllostachyspubescens* forest in subtropical China. SAR had no significant effects on both soil respiration rate and soil microbial biomass. The soil bacterial diversity analysis indicated that only T3 treatment (pH 2.5) showed a significant effect on soil bacterial diversity relative to that of CK, and the higher acid load (T2 and T3) increased the soil bacterial diversity in terms of Shannon index. Overall, the impacts of acid rain in bamboo forest ecosystem was highlighted in this study and studying soil respiration and microbial community in bamboo forest (subtropical forest) subject to SAR treatments could contribute for understanding their functions in C cycling/ecosystem C flux.

## Acknowledgments

This work was supported by Zhejiang Provincial Natural Science Foundation of China (LY15C160005 and LQ19C160013) and Foundation of Zhejiang Educational Committee (Y201329652).

## Author Contributions

Nan Wang performed the experiments and wrote the paper; Xiaocheng Pan designed the experiments and reviewed the paper.

## Conflicts of Interest

The authors declare no conflict of interest.

